# Expanding the toolkit for ploidy manipulation in *Chlamydomonas reinhardtii*

**DOI:** 10.1101/2024.10.01.616035

**Authors:** Antoine Van de Vloet, Lucas Prost-Boxoen, Quinten Bafort, Yunn Thet Paing, Griet Casteleyn, Lucile Jomat, Stéphane D. Lemaire, Olivier De Clerck, Yves Van de Peer

**Affiliations:** VIB-UGent Center for Plant Systems Biology, Ghent, Belgium; Department of Plant Biotechnology and Bioinformatics, Ghent University, Ghent, Belgium; Department of Biology, Ghent University, Ghent, Belgium; Laboratoire de Biologie Computationnelle et Quantitative, Institut de Biologie Paris-Seine, UMR 7238, Sorbonne Université, CNRS, Paris, France; Biofonderie de l’Alliance Sorbonne Université, UAR2037, Sorbonne Université, CNRS, Université de Technologie de Compiègne, Paris, France; College of Horticulture, Academy for Advanced Interdisciplinary Studies, Nanjing Agricultural University, Nanjing, China; Department of Biochemistry, Genetics and Microbiology, University of Pretoria, Pretoria, South Africa

**Author notes:** These authors contributed equally to this work (Antoine Van de Vloet, Lucas Prost-Boxoen).

**Keywords:** *Chlamydomonas reinhardtii*, experimental model, plant evolution, synthetic ploidy, polyploidy, transformation, mating, flow cytometry

## Abstract

**Summary:**

- Whole genome duplications, widely observed in plant lineages, have significant evolutionary and ecological impacts. Yet, our current understanding of the direct implications of ploidy shifts on short- and long-term plant evolution remains fragmentary, necessitating further investigations across multiple ploidy levels.
- *Chlamydomonas reinhardtii*, a haploid green alga, is a valuable model organism with profound potential to study the impact of ploidy increase on the longer-term in a laboratory environment. This is partly due to the ability to successfully increase the ploidy level.
- We developed a strategy to engineer ploidy in *Chlamydomonas reinhardtii* using a collection of non-interfering antibiotic selectable markers. This approach allows to induce higher ploidy levels in *Chlamydomonas reinhardtii* and is applicable to field isolates, which expands beyond specific auxotroph laboratory strains and broadens the genetic diversity of parental haploid strains that can be crossed. We implement flow cytometry for precise measurement of the genome size of strains of different ploidy.
- We demonstrate the creation of diploids, triploids and tetraploids by engineering North American field isolates, broadening the application of synthetic biology principles in *Chlamydomonas reinhardtii*.
- Our study greatly facilitates the application of *Chlamydomonas reinhardtii* to study polyploidy, both in fundamental and applied settings.

## Introduction

Polyploidy, the heritable condition of having more than two complete sets of chromosomes resulting from a whole genome duplication (WGD), is widespread across the tree of life, particularly in flowering plants (Otto & Whitton, 2000; Soltis *et al*., 2015; Van de Peer *et al*., 2017). Although WGD is often considered as an important evolutionary driver within the flowering plants, the initial cost of ploidy increase is substantial (Comai, 2005; Hollister, 2015; Yant & Bomblies, 2015; Baduel *et al*., 2018; Mortier *et al*., 2024). Many of the immediate morphological, developmental, phenological, and physiological effects of polyploidization in plants have been reported (as reviewed by for instance Doyle & Coate, 2019; Bomblies, 2020; Clo & Kolář, 2021), and since many of these immediate effects present challenges rather than opportunities, it is still unclear to what extent they contribute to longer-term evolutionary benefits and polyploid establishment (Soltis *et al*., 2015; Dodsworth *et al*., 2016; Baduel *et al*., 2018; Li *et al*., 2021; Wang *et al*., 2021). Currently, most knowledge on the longer-term evolutionary effects of polyploidy is derived from studies on naturally occurring (paleo)polyploids (Bomblies, 2020), and experimental data covering many generations after WGD is scarce. To gain further insights into the mechanisms underlying both the short- and long-term effects of polyploidy, evolutionary experiments using artificial polyploids and their lower-ploidy ancestors are needed. To this end, we need model systems in which polyploidy can be readily engineered, that are easily manipulated and replicated, and preferably have short generation times facilitating the study of many generations in a relatively short time span.

The green microalga *Chlamydomonas reinhardtii* (hereafter referred to as Chlamydomonas) is a well-established model organism and offers a valuable model for research on polyploidy and evolution. It provides the experimental tractability of other microscopic model systems such as yeast, while being phylogenetically closer to angiosperms (Bafort *et al*., 2023). Chlamydomonas is also a synthetic biology chassis amenable to advanced genome engineering (Crozet *et al*., 2018; Vavitsas *et al*., 2019; Goold *et al*., 2024) providing an ideal platform for engineering synthetic ploidy. Although most research has been carried out with a limited set of inbred laboratory strains, population genomic analyses on field isolates demonstrate a high nucleotide diversity in the species (Craig *et al*., 2019). Chlamydomonas has a haplontic life cycle and only enters a diploid phase during sexual reproduction, forming a dormant zygospore (Harris, 2009). A variable minority of gametic fusion products fails to form zygospores, resulting in mitotic growth of diploid cell lines (Ebersold, 1963, 1967; Kariyawasam *et al*., 2019). Leveraging this phenomenon, Ebersold (1967) developed a method for isolating diploid Chlamydomonas colonies on selective media, through complementation of recessive auxotrophic mutations. Using this approach, past efforts have successfully produced diploids (Ebersold, 1967; Sosa *et al*., 1978; Palombella & Dutcher, 1998; Bellafiore *et al*., 2002) and even triploid and tetraploid strains (Matagne & Mathieu, 1983; Matagne & Yu, 1987; Kariyawasam *et al*., 2019).

However, the current methodology relies on auxotrophic nuclear recessive mutations obtained through non-specific mutagenesis which has limited the application of the crosses to specific laboratory strains, excluding the genetically diverse set of field isolates. Moreover, these mutants may suffer substantially from off-target mutations, while currently more precise transformation methods have been developed for Chlamydomonas (Crozet *et al*., 2018). Previous studies verified the ploidy level of polyploid Chlamydomonas through morphological, biochemical or genetic means (Harris, 2009). Morphological evidence often involves chromosome counts or differences in cell size (e.g. Ebersold, 1967; Matagne & Yu, 1987), while biochemical verification quantifies DNA content relative to cell number using DNA-specific stains (Stein-Taylor, 1973; Valle *et al*., 1981; Zachleder, 1984; Kariyawasam *et al*., 2019). Genetic proof can be obtained through specific crosses (Gillham, 1969), or a PCR-based assay for mating type (Werner & Mergenhagen, 1998; Zamora *et al*., 2004). Flow cytometry of individual nuclei stained with DNA-specific stains has recently been optimized for the genus *Chlamydomonas* (Čertnerová, 2021), allowing for precise genome size quantification without biases such as differences in cell concentration or extraction efficiency. It also offers a higher resolution, with the potential to detect aneuploidies (Sliwinska *et al*., 2022).

Here we present a synthetic biology approach for inducing and measuring polyploidy in *Chlamydomonas reinhardtii*, leveraging a collection of antibiotic selectable markers and flow cytometry. Our strategy builds on Ebersold’s complementation principle, using haploid strains engineered with genetic circuits conferring resistance to diverse antibiotics. This enables the use of any pair of mating-compatible parental strains including field isolates. Flow cytometry was applied to quantify synthetic ploidy levels, offering high-resolution measurements of genome size. Our results confirm that this strategy, using field isolates from the North American continent, successfully yields diploid, triploid, and tetraploid Chlamydomonas strains. This work provides a methodology that not only induces polyploidy in diverse strains but also accurately quantifies genome size.

## Materials and Methods

### Strains, their maintenance and mating type verification

We selected three pairs of haploid Chlamydomonas strains with opposite mating types. The pairs were chosen to ensure that their pairwise genetic distances span a broad spectrum, based on nucleotide diversity data published in Craig *et al*. (2019), to allow the possibility of including different levels of heterozygosity in our potential polyploid progeny. We selected four North American field isolates, CC-2931 (*mt-*), CC-2932 (*mt+*), CC-2935 (*mt-*) and CC-2937 (*mt+*) (Chlamydomonas Resource Center; Lefebvre *et al*., 2019), and two Japanese field isolates, NIES-2464 (*mt-*) and NIES-2463 (*mt+*) (National Institute for Environmental Studies, NIES; Nakada *et al*., 2010). When not in active cultivation, strains were preserved on Tris-acetate-phosphate (TAP) (Gorman & Levine, 1965) agar plates under low light conditions (50 μmol·m^− 2·^s^−1^ PPFD) at 18°C. Before initiating crosses, mating types were confirmed by Chelex DNA extraction (Cao *et al*., 2009) followed by polymerase chain reaction (PCR) targeting the mating type locus (Zamora *et al*., 2004).

### Crossing strategy

A schematic representation of the crossing strategy is shown in Figure 1.a. The *mt-* strain of each selected haploid pair was engineered with a genetic circuit conferring antibiotic resistance to marker A, while the *mt+* strains were engineered three different times with circuits for resistance markers B, C, or D. The *mt-*/A haploid was crossed with the *mt+*/B haploid and the progeny was grown on double selective medium (compounds A and B present). Following Ebersold’s approach (Ebersold, 1967), diploids resistant to both selectable markers were isolated. These A/B-resistant diploids (*mt-*) were then crossed with the C-resistant *mt+* strain. Selection on triple selective medium allowed for isolation of triploid strains resistant to antibiotics A, B, and C. Finally, this triploid strain (*mt-*) was crossed with the D-resistant *mt+* strain to generate tetraploid strains. A list of all transformed haploids used to test this crossing scheme is provided in Table 1.

**Figure 1.**
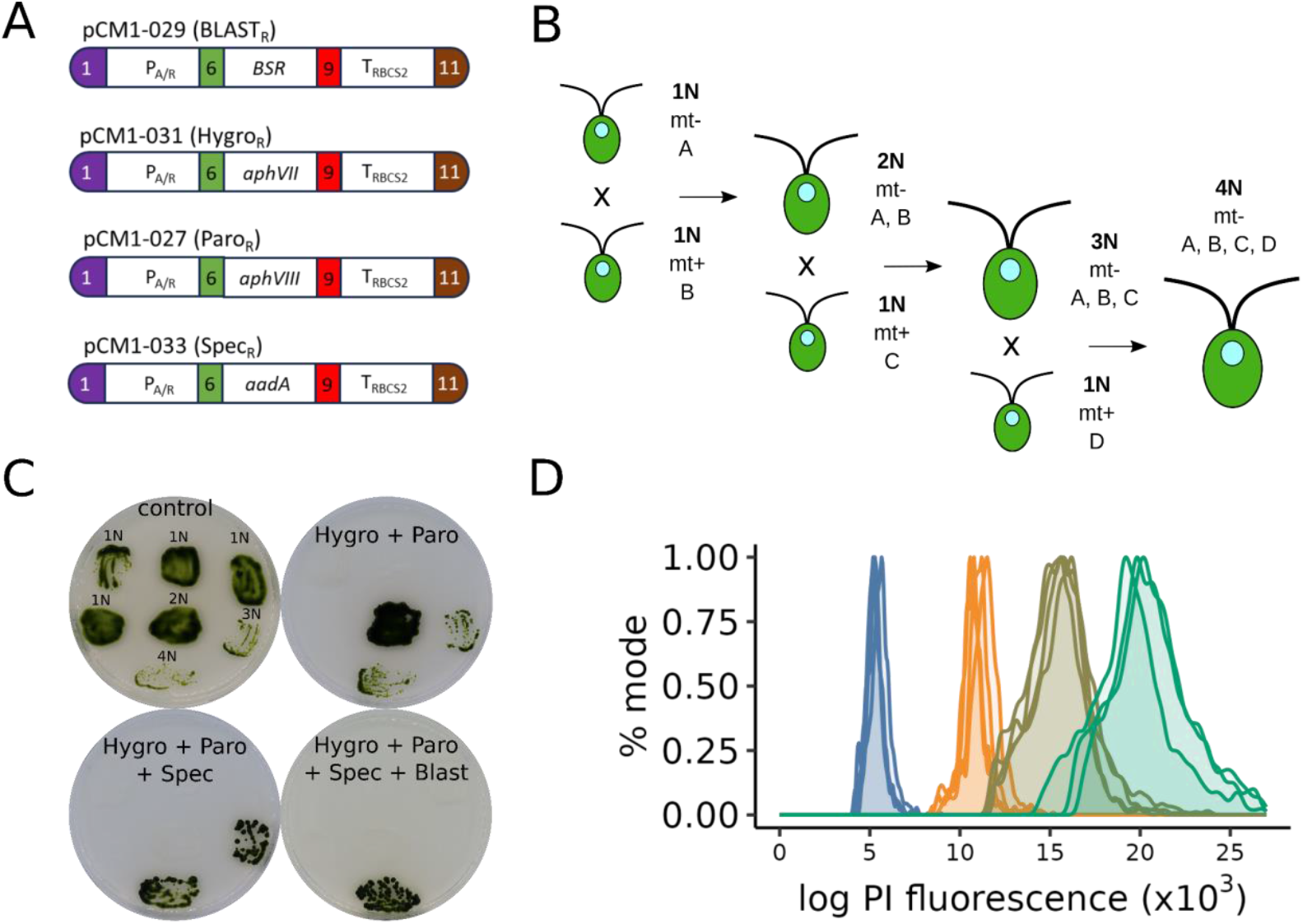
Generation and analysis of polyploid Chlamydomonas offspring. **A:** Genetic circuits enabling synthetic ploidy approaches. The genetic circuits conferring resistance to blasticidin S (pCM1-029), hygromycin (pCM1-031), paromomycin (pCM1-027), spectinomycin (pCM1-033) comprise a constitutive promoter (PA/R + 5′UTR of RBCS2), the resistance coding sequence and a terminator (3′UTR of RBCS2 + TRBCS2) and were assembled using Golden Gate cloning (De Carpentier *et al*., 2020). The numbers 1, 6, 9, and 11 stand for the standard fusion sites used for cloning within the standard of the Chlamydomonas Modular Cloning toolkit (Crozet *et al*., 2018). **B:** Schematic representation of the crossing strategy used to obtain tetraploid Chlamydomonas starting from minus (*mt-*) and plus (*mt+*) haploid transformant strains. Capital letters represent unique antibiotic resistance markers inserted into the haploid genomes. **C**: Growth on agar plates of selected progeny. The growth and resistance is shown for haploid parents and diploid, triploid and tetraploid progeny, on both a control plate (TAP medium without antibiotics, top left plate) and selective plates with two (20 mg.L^-1^ hygromycin B and 20 mg.L^-1^ paromomycin, top right plate), three (an additional 100 mg.L^-1^ spectinomycin, bottom left plate) and four (an additional 50 mg.L^-1^ blasticidin S, bottom right plate) antibiotics added. **D:** Flow cytometry profiles of PI-stained nuclei. Haploids are shown in blue, diploids in orange, triploids in khaki and tetraploids in green peaks.

**Table 1.** Overview of the transformed haploid strains used as input to test the proposed crossing scheme. Given is: the strain ID within the collection that maintains this strain, the confirmed mating type, the genetic circuit that was introduced in the genome to confer resistance to a particular antibiotic substance (insert), the antibiotic substance to which that specific insert confers resistance, the minimum effective concentration used to implement the selective environment and the reference we use in this study for that transformant strain.

| Strain ID | Mating type | Insert | Antibiotic resistance | Concentration (mg/L) | Reference |
| --- | --- | --- | --- | --- | --- |
| CC-2935 | mt- | A | Hygromycin B | 20 | H1 |
| CC-2937 | mt+ | B | Paromomycin | 20 | H2 |
| CC-2937 | mt+ | C | Spectinomycin | 100 | H3 |
| CC-2937 | mt+ | D | Blasticidin S | 50 | H4 |
| CC-2931 | mt- | A | Hygromycin B | 20 | H5 |
| CC-2932 | mt+ | B | Paromomycin | 20 | H6 |
| CC-2932 | mt+ | C | Spectinomycin | 100 | H7 |
| CC-2932 | mt+ | D | Blasticidin S | 50 | H8 |
| NIES-2464 | mt- | A | Hygromycin B | 20 | H9 |
| NIES-2463 | mt+ | B | Paromomycin | 20 | H10 |
| NIES-2463 | mt+ | C | Spectinomycin | 100 | H11 |
| NIES-2463 | mt+ | D | Blasticidin S | 50 | H12 |

### Transformations

We chose four selectable markers known to respectively confer resistance of Chlamydomonas to hygromycin, paromomycin, spectinomycin and blasticidin S. These antibiotics were selected because they act at the post-translational level, ensuring that the compound does not induce unwanted changes in the DNA sequence of the progeny, and because these markers do not suffer from interference in their mode of action (De Carpentier *et al*., 2020). The constructs used to engineer Chlamydomonas were generated using the Chlamydomonas modular cloning (MoClo) toolkit as described in Crozet *et al*. (2018). The genetic circuits were built into level 1 plasmids from level 0 parts to enable antibiotic resistance under the control of the PA/R promoter coupled to the 5′UTR of RBCS2 (pCM0-020) and the 3′UTR of RBCS2 coupled to terminator TRBCS2 (pCM0-115) (Schroda *et al*., 2002; De Carpentier *et al*., 2020). The level 1 plasmids used for Chlamydomonas engineering were pCM1-031 for hygromycin (pCM0-073), pCM1-027 for paromomycin (pCM0-074), pCM1-033 for spectinomycin (pCM-076) and pCM1-029 for blasticidin S (pCM0-120) (Figure 1.a), as described and characterized in De Carpentier *et al*., 2020. Chlamydomonas cells were grown in TAP medium to early exponential phase (∼1–2 × 106 cells.mL-1), and then concentrated 100 times in TAP + 60 mM sucrose. After incubating 250 µL of cells with DNA (55 fmol of purified genetic circuit excised with BbsI-HF; New England Biolabs) on ice for 20 min, the cells were electroporated (2000 V.cm-1, 25 µF, no shunt resistance) and incubated for 16–20 hours in 10 mL of TAP + 60 mM sucrose prior to be plated on TAP-agar plates supplemented with appropriate antibiotic(s). Transformants were selected on TAP-agar medium containing hygromycin B (20 mg.L-1), paromomycin (20 mg.L-1), spectinomycin (100 mg.L-1) or blasticidin S (50 mg.L-1). Colonies appeared after 5–7 days of growth in continuous light (50 µmol.m−2.s−1 PPFD) at 25°C.

### Gametogenesis and mating

Strains of opposite mating type were cultivated in 5 mL of TAP medium in Pyrex Erlenmeyer Flasks at 25°C with ∼200-250 µmol.m^− 2^.s^− 1^ PPFD under agitation at 175 rpm. After 3-4 days, cultures reached stationary phase (∼10^7^ cells.mL^-1^). The cells were centrifuged at 2000 g for 5 min, and the resulting cell pellets were resuspended in 7 mL of TAP-N medium, wherein NH_4_Cl is substituted with KCl to remove the nitrogen source to induce gametogenesis. The cultures were then incubated overnight under low light (∼50 µmol.m^− 2^.s^− 1^ PPFD) and agitation at 75 rpm to promote gametogenesis. Gamete presence was verified by the active movement of the cells using an inverted light microscope (Tulin, 2019). To initiate mating, samples of 1.5 mL from both the *mt+* and *mt-* gamete cultures were mixed and incubated under low light without agitation for 2 hours. Afterwards, 1.5 mL aliquots were plated on the appropriate selective TAP agar plates and incubated under low light conditions without agitation, with daily monitoring for colony appearance. After 24 hours, thick-walled zygospores were observed on the plate under a microscope, indicating successful mating.

### Selection and isolation of candidate colonies

Macroscopic colonies became visible on selective plates after about 5 days for diploids and 8-10 days for triploids and tetraploids. Early appearing colonies, particularly those observed before zygote germination, were identified as candidate colonies. These candidates were picked up and streaked on a fresh plate with identical antibiotic concentrations as the selective plate. After a 10-day incubation under low light (∼50 µmol.m^− 2^.s^− 1^ PPFD), colony growth was assessed to confirm resistance to all antibiotics in the medium.

### Synchronizing conditions

Cultures were synchronized in 5 mL TAP medium, undergoing 12h:12h light:dark cycles with continuous agitation at 175 rpm. These cycles included a phase with intense light (∼220 µmol.m^− 2^.s^− 1^ PPFD) at 28°C followed by a complete dark phase at 18°C. These conditions were maintained for 15 days, incorporating regular bottlenecks for optimal synchronization.

### Genome size estimation

The genome size of a culture was measured using flow cytometry on nuclei stained with propidium iodide (PI). This protocol is based on protocol 6 in Čertnerová (2021). A 2 mL sample from a mid-exponential synchronized culture was harvested 2 hours post-initiation of the light-warm phase, targeting early G1 cells. This sample was centrifuged at 6000 g for 7 minutes, after which the supernatant was removed. Approximately 10 glass beads of 1.5 mm diameter were added to a 2 mL Eppendorf tube containing 550 μL of ice-cold lysis buffer LB01 (Dpooležel *et al*., 1989) and the cell pellet. The cells were disrupted for 3 min at 25 Hz using a Retsch MM400 mixer mill. The resultant sample was filtered through a 42 μm nylon mesh. This was followed by an incubation period at 4°C of approximately one hour, during which the sample spontaneously split into two phases: a lower green layer containing remaining cell debris and pigments, and a colorless upper layer in which the nuclei remain in solution. 200 μL of this upper layer was isolated and added to a staining solution composed of 550 μL of LB01 lysis buffer, 50 μg.mL^− 1^ PI, 50 μg.mL^− 1^ RNase IIA and 2 μL.mL^− 1^ β-mercaptoethanol. Following a 10-minute dark incubation at 4°C, the stained samples were analyzed with an Attune NxT flow cytometer (ThermoFisher Scientific, Waltham). Nuclei were identified by using a yellow excitation laser (561 nm, 50 mW, emission filter 620/15, channel YL2) and subsequently plotting side scatter area (SSC-A) against PI intensity (Figure 2.a). When the nuclei were gated, only the G1 populations were further studied. The median value of the distribution of all intensities in this nuclei population (Median Nuclei Value, MNV) was further used as a proxy for genome size and ploidy level (Figure 2.b). Imaging flow cytometry was used to verify that we correctly identified nuclei from all other measured particles (Figure 2.c). These images were generated using a BD Biosciences FACS imaging-enabled prototype cell sorter that is equipped with an optical module allowing multicolor fluorescence imaging of fast-flowing cells in a stream, enabled by BD CellView^TM^ Image Technology based on fluorescence imaging using radiofrequency-tagged emission (FIRE) (Diebold *et al*., 2013; Schraivogel *et al*., 2022).

**Figure 2.**
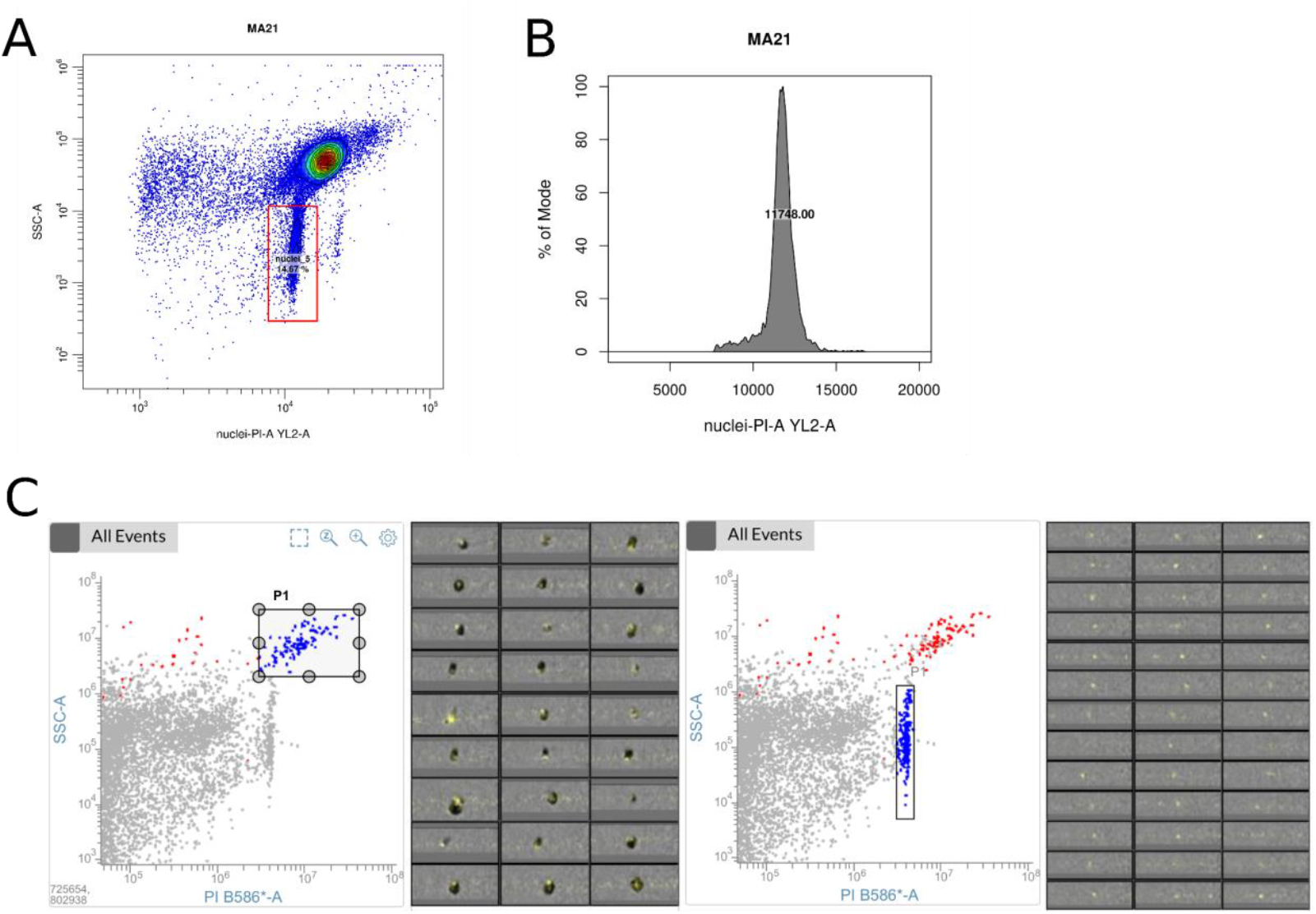
Gating strategy for PI-stained nuclei. **A:** PI-stained nuclei appear as a separate population by plotting side scatter of the particles (SSC-A) over their PI intensity value (nuclei PI-A YL2-A). **B:** The median value of the distribution of PI intensity values of the gated nuclei is the Median Nuclei Value (MNV) and is used as a proxy for genome size of a culture. **C:** Imaging data on the different populations in our SSC-A vs. PI signal plot. Cells with illuminated, stained nuclei in yellow (left) are well separated from stained, isolated nuclei (right). Other events in this plot were confirmed as noise such as cell debris.

## Results

Successful growth of the progeny on the appropriate selective medium was used as the first indicator of candidate polyploids (Figure 1.b) and flow cytometry was subsequently applied to confirm the genome size (Figure 1.c). Flow cytometry profiles for haploid strains showed a median log PI fluorescence value around 5500, and the profiles for higher ploidy levels scaled linearly with the ploidy level. Although the cell concentrations in our gamete and mating cultures were of the same order of magnitude, the number of colonies that grew on selective media decreased over ploidy levels (Table 2). Additionally, we identified several false positive colonies which grew on the selective medium but did not show increase in ploidy based on flow cytometry analysis (Table 2). The overall success rate seems highest for creating diploids and generally lowest for creating triploids.

**Table 2.** Overview of all the successful crosses obtained in this study. Given is: the reference that we give the progeny of this cross in this study, information on both parental strains (reference, mating type, resistance), the resistance of the progeny confirmed by selective growth, the mating type phenotype of the progeny, the number of days the initial colony grew on resistance plates before it was considered viable and was picked up, the amount of colonies tested with flow cytometry is indicated, and the amount of colonies that tested positive for a ploidy increase. Finally, the intended Relative Genome Size (RGS, calculated as the median PI fluorescence for that strain divided by the sum of median fluorescence values of all haploids used in the crossing) is indicated, together with the initial measured RGS of one of the successful ploidy increase events (measured after the days of growth indicated in the table + two weeks of synchronization in liquid culture post spawn of the progeny).

| Reference | P1 | mt<br>P1 | AB<br>resistance<br>P1 | P2 | mt P2 | AB resistance<br>P2 | AB resistance<br>progeny | Days of<br>growth | Tested<br>colonies | Positive<br>colonies | Measured<br>RGS | Intended<br>RGS |
| --- | --- | --- | --- | --- | --- | --- | --- | --- | --- | --- | --- | --- |
| D1 | H1 | mt- | Hygromycin<br>(H) | H6 | mt+ | Paromomycin<br>(P) | H, P | 5 | 25 | 25 | 1.919403 | 2 |
| D2 | H1 | mt- | Hygromycin<br>(H) | H2 | mt+ | Paromomycin<br>(P) | H, P | 5 | 25 | 25 | 2.063912 | 2 |
| D3 | H5 | mt- | Hygromycin<br>(H) | H6 | mt+ | Paromomycin<br>(P) | H, P | 5 | 25 | 25 | 2.021283 | 2 |
| D4 | H5 | mt- | Hygromycin<br>(H) | H2 | mt+ | Paromomycin<br>(P) | H, P | 5 | 25 | 25 | 2.036123 | 2 |
| T1 | D1 | mt- | H, P | H7 | mt+ | Spectinomycin<br>(S) | H, P, S | 8-10 | 6 | 5 | 2.828334 | 3 |
| T2 | D1 | mt- | H, P | H3 | mt+ | Spectinomycin<br>(S) | H, P, S | 8-10 | 9 | 6 | 2.938732 | 3 |
| T3 | D2 | mt- | H, P | H7 | mt+ | Spectinomycin<br>(S) | H, P, S | 8-10 | 7 | 7 | 2.601228 | 3 |
| T4 | D2 | mt- | H, P | H3 | mt+ | Spectinomycin<br>(S) | H, P, S | 8-10 | 8 | 1 | 2.510176 | 3 |
| T5 | D3 | mt- | H, P | H7 | mt+ | Spectinomycin<br>(S) | H, P, S | 8-10 | 9 | 1 | 2.639808 | 3 |
| T6 | D3 | mt- | H, P | H3 | mt+ | Spectinomycin (S) | H, P, S | 8-10 | 9 | 8 | 2.769077 | 3 |
| T7 | D4 | mt- | H, P | H7 | mt+ | Spectinomycin (S) | H, P, S | 8-10 | 7 | 7 | 2.854335 | 3 |
| T8 | D4 | mt- | H, P | H3 | mt+ | Spectinomycin (S) | H, P, S | 8-10 | 9 | 6 | 2.884225 | 3 |
| Q1 | T2 | mt- | H, P, S | H8 | mt+ | Blasticidin (B) | H, P, S, B | 8-10 | 7 | 5 | 3.54116 | 4 |
| Q2 | T2 | mt- | H, P, S | H4 | mt+ | Blasticidin (B) | H, P, S, B | 8-10 | 3 | 3 | 3.252065 | 4 |
| Q3 | T3 | mt- | H, P, S | H8 | mt+ | Blasticidin (B) | H, P, S, B | 8-10 | 3 | 2 | 3.344298 | 4 |
| Q4 | T4 | mt- | H, P, S | H8 | mt+ | Blasticidin (B) | H, P, S, B | 8-10 | 2 | 2 | 3.617856 | 4 |
| Q5 | T6 | mt- | H, P, S | H8 | mt+ | Blasticidin (B) | H, P, S, B | 8-10 | 6 | 4 | 3.359455 | 4 |
| Q6 | T6 | mt- | H, P, S | H4 | mt+ | Blasticidin (B) | H, P, S, B | 8-10 | 4 | 4 | 3.338466 | 4 |
| Q7 | T8 | mt- | H, P, S | H8 | mt+ | Blasticidin (B) | H, P, S, B | 8-10 | 3 | 3 | 3.752124 | 4 |
| Q8 | T8 | mt- | H, P, S | H4 | mt+ | Blasticidin (B) | H, P, S, B | 8-10 | 2 | 2 | 3.433374 | 4 |

This methodology was systematically applied to all six selected strains, initially yielding 12 unique successfully transformed haploids that were used as input (Table 1). All possible combinations at each step of the crossing scheme were tested with these 12 transformed haploids. Table 2 lists all successful crosses, defined as those resulting in the intended ploidy increase being present among the progeny. Notably, none of the crosses with the Japanese (NIES) strains succeeded. At the first step and for all crosses involving at least one Japanese strain, growth on double selective medium was minimal, and the few colonies that did form proved haploid using flow cytometry. We observed a low probability of successful gametogenesis in these haploid mating cultures. Microscopic observation of Japanese gamete cultures revealed a lethargic state, with cells present in (often large) aggregates (Figure S1), instead of the typical vigorously moving individual gametes (Tulin, 2019). This was followed by the absence of zygotes after mating.

## Discussion

Ever since Ebersold (1967) developed a method to create vegetative diploids using complementation, ploidy increase in Chlamydomonas has been part of the toolkit of the Chlamy-geneticist (Ebersold, 1967; Loppes, 1969; Matagne & Orbans, 1980; Matagne & Mathieu, 1983; Matagne & Yu, 1987; Palombella & Dutcher, 1998; Bellafiore *et al*., 2002; Kariyawasam *et al*., 2019; Bafort *et al*., 2023). However, Ebersold’s double complementation method relies on auxotrophic mutations, which limits its application to specific auxotrophic mutant Chlamydomonas strains. We expanded this methodology by proposing a transformation-based synthetic biology approach that enables the combination of any mating-compatible genomes, including genetically diverse field isolates. This new method offers broader applicability and the capability to produce diploid, triploid, and tetraploid strains. Despite the expected versatility of our method, we encountered specific challenges with strains from the NIES collection. Initially, we had no reason to anticipate these issues, as previous research indicated that Japanese strains can form viable hybrids with North American strains (Nakada *et al*., 2014). However, our microscopic observations suggest that gametogenesis failure occurred in the Japanese mating cultures. The absence of zygotes and vegetative diploids in these crosses, and the fact that crosses between the Japanese strains also failed, supports the idea that the problem lies with gametogenesis in these strains rather than with mating-incompatibilities.

We observed a negative trend between the success rate of ploidy increase and the level of ploidy. Specifically, triploid and tetraploid colony formation was less frequent compared to formation of diploid colonies. We are cautious in trying to explain this increasing false positive rate as various causes may underlie this phenomenon. One possible explanation is a reduced efficiency in forming functional gametes, or the delayed or slowed growth of the resulting polyploids, allowing the appearance of vegetative polyploids to coincide with germination of the zygotes. This delay could allow vegetative polyploid cells to appear concurrently with zygote germination. Consistently, our data indicate that colony formation occurred progressively later with increasing ploidy levels, as observed in previous studies (Matagne & Yu, 1987).

This study pioneers the progressive creation of diploid, triploid and tetraploid strains of Chlamydomonas, validated by flow cytometry-based genome size quantification, a technique that has become a gold standard for genome size estimation in plant science (Sliwinska *et al*., 2022). Our results show that we extracted nuclei in a reliable way and could accurately quantify genome sizes in our polyploid progeny. Flow cytometry analysis provides a more precise and reliable estimate of genome size in Chlamydomonas compared to previous studies in this context. Traditionally there was a strong reliance on phenotypic markers, such as rescue of wild type phenotypes on selective media, chromosome counts, or an increase in cell size as proxy (e.g. Ebersold, 1967; Matagne & Yu, 1987). Although suggestive of ploidy elevation, these data may reflect other causes such as meiotic recombinants, aneuploidies or strain effects. More direct efforts in quantifying nuclear DNA content of polyploid Chlamydomonas products were based on DNA-specific fluorometric data (e.g. Kariyawasam et al., 2019). Although standardized by the number of harvested cells, this approach may suffer from substantial variability in sample extraction efficiency or noise on the cell count estimates. The flow cytometry method we apply here measures a DNA-specific fluorescence signal in individual nuclei. With the implementation of control samples, this allows for a precise estimate of increase in genome size and the different ploidies present in a culture.

## Conclusion & prospects

The green alga *Chlamydomonas reinhardtii*, a single celled plant that is often referred to as ‘the green yeast’ (Goodenough, 1992), is a widely used model organism that had a pivotal role in advancing our understanding of essential biological processes such as photosynthesis, structural biology, cell cycle regulation and more (Sasso *et al*., 2018). In addition, its fully sequenced genome (Craig *et al*., 2023), short generation time and ease of culturing (Harris, 2001) make it a particularly suitable system for studying evolution in a controlled laboratory environment. Together with the existing methodology to increase its ploidy level, it has been proposed as an excellent model to study whole genome duplications and their evolutionary consequences (Bafort *et al*., 2023). Our work significantly expands the toolkit of ploidy increase in Chlamydomonas. By leveraging genetic circuits that confer antibiotic resistance, we have developed a method that enables generating higher-order ploidy. Importantly, the methodology does not rely on the use of auxotrophic mutants, thereby broadening its applicability beyond traditional laboratory strains. Future research can explore the auto-allopolyploid spectrum by elegantly experimenting with varying degrees of heterozygosity between haploid sub genomes, given the substantial nucleotide diversity in the species (Craig *et al*., 2019). The proposed crossing scheme allows us to reach at least the tetraploid level, but could in theory be extended further, provided there are sufficient selectable markers that do not interfere with one another. Additionally, the implementation of flow cytometry for genome size quantification provides not only an accurate means of confirming ploidy increase, but also offers the potential to detect large scale genome size alterations following WGD, such as aneuploidies (Matagne & Orbans, 1980). This aspect opens new avenues for exploring genome stability in the aftermath of polyploidy and could yield insights into the mechanisms by which polyploid genomes evolve over time.

Our synthetic biology approach establishes a foundation for future exploration of synthetic ploidy in Chlamydomonas, enabling investigation into both immediate and long-term effects of polyploidy and genome duplications for both fundamental research and biotechnological applications. Beyond its fundamental interest, notably in the field of evolutionary biology, our method of genome merging might also be of biotechnological interest. As industries transition toward more sustainable production methods, microalgae bioproduction emerges as a critical technology, capable of reducing resource use and minimizing carbon footprints. Microalgae are regarded as promising synthetic biology chassis, capable of producing valuable compounds through their natural ability to fix CO2 in a sunlight-driven and sustainable process. Our work opens the door for exploiting hybrid vigor and other beneficial effects of genome merging such as increased growth rates, yield or stress resistance (Radakovits *et al*., 2010; Kwak *et al*., 2017; Goold *et al*., 2024). These advancements could significantly enhance the biotechnological applications of algal bioproduction, particularly in the context of climate change mitigation and renewable bioproduct development.

## Acknowledgements

YVdP acknowledges funding from the European Research Council (ERC) under the European Union’s Horizon 2020 research and innovation program (No. 833522). ODC is supported by EMBRC Belgium and the Research Foundation Flanders (I001621N). AVDV and YVdP acknowledge funding from Ghent University (Methusalem funding, BOF.MET.2021.0005.01). LP-B and QB were awarded PhD scholarships by the Fonds Wetenschappelijk Onderzoek (FWO) of Flanders (Grant Nos. 11H0426N and 1168420N, respectively). SDL and LJ acknowledge funding from Sorbonne Université and Région Ile de France (DIM BioconvS and SESAME Filières BFB). The authors thank Dr. Eylem Aydogdu for helping set up the Chlamydomonas system and Gert Van Isterdael, Elien Ruyssinck and Julie Van Duyse from the VIB Flow Core, VIB Center for Inflammation Research, Ghent, Belgium, for their insights and help on the flow cytometry.

## Author contributions

**AVdV, LPB, QB, ODC and YVdP** designed the study. **SDL and LJ** carried out the transformations. **AVdV, GC and YTP** carried out the strain maintenance, crosses and the flow cytometry. **AVdV and LPB** wrote the original draft. **QB, SDL, LJ, ODC and YVdP** edited the manuscript.

## Data availability

All haploid transformant strains will be made available through the Chlamydomonas Resource Center (https://www.chlamycollection.org/, Lefebvre *et al*., 2019).

## Supporting Information

**Figure S1.**
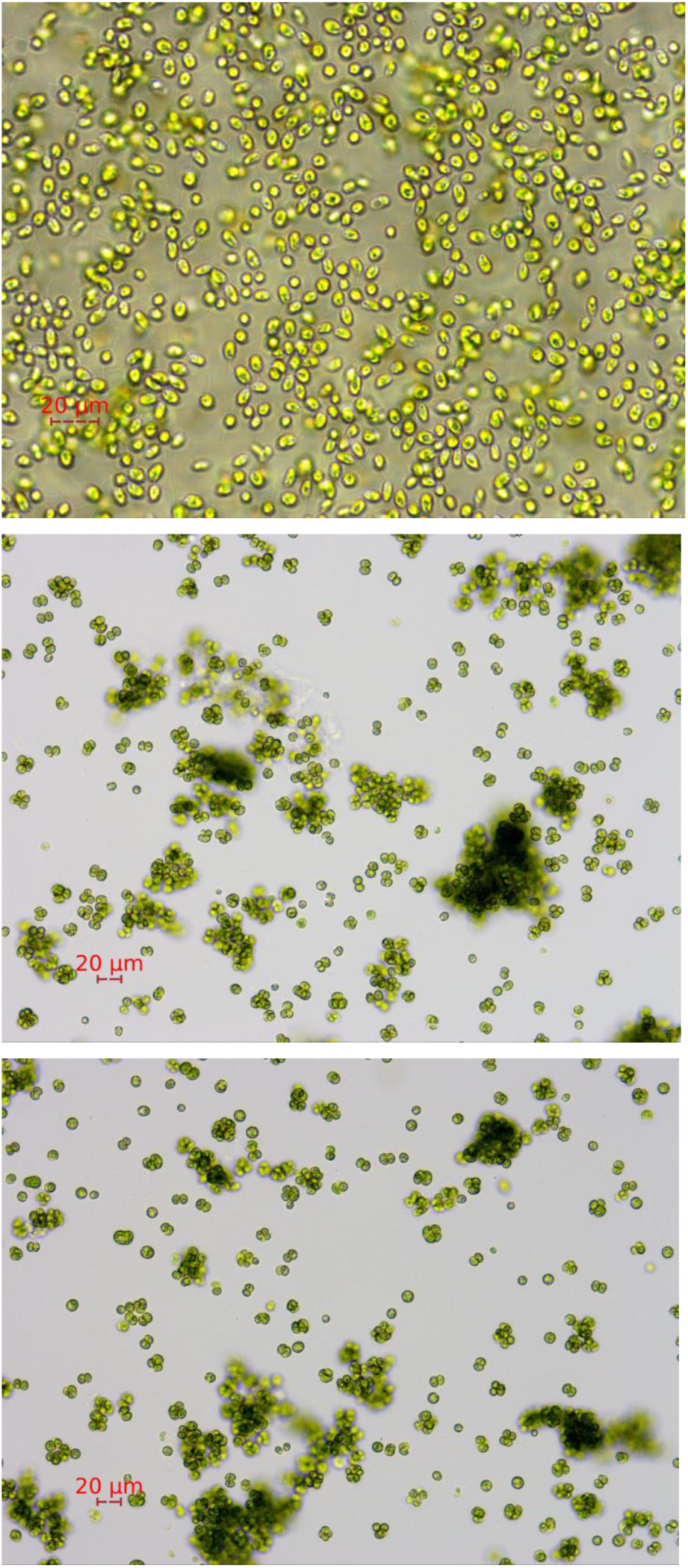
Microscopic images of gamete cultures. Pictures were taken with a ToupTek ToupCam camera mounted onto a Olympus CKX53 inverted microscope. **Top:** gametes of H6 (cc-2932 ParoR). This culture displays individual moving cells with both cilia present. Magnification Ph 40×/0,55. **Middle:** failed gametogenesis in H9 (NIES-2464 HygroR). Cells appear in aggregates without movement. Magnification Ph 20×/0,4. **Bottom:** failed gametogenesis in H10 (NIES-2463 ParoR). Cells appear in aggregates without movement. Magnification Ph 20×/0,4.

